# Individual similarities and differences in eye-movement-related eardrum oscillations (EMREOs)

**DOI:** 10.1101/2023.03.09.531896

**Authors:** Cynthia D King, Stephanie N Lovich, David LK Murphy, Rachel Landrum, David Kaylie, Christopher A Shera, Jennifer M Groh

**Author notes:** Department of Psychiatry and Behavioral Sciences, Duke University (David LK Murphy). Louisiana State University School of Medicine.

## Abstract

We recently discovered a unique type of low-frequency otoacoustic emission (OAE) time-locked to the onset (and offset) of saccadic eye movements and occurring in the absence of external sound (Gruters et al., 2018). How and why these eye-movement-related eardrum oscillations (EMREOs) are generated is unknown, with a role in visual-auditory integration being the likeliest candidate. Clues to both the drivers of EMREOs and their purpose can be gleaned by examining responses in normal hearing human subjects. Do EMREOs occur in all individuals with normal hearing? If so, what components of the response occur most consistently? Understanding which attributes of EMREOs are similar across participants and which show more variability will provide the groundwork for future comparisons with individuals with hearing abnormalities affecting the ear’s various motor components. Here we report that in subjects with normal hearing thresholds and normal middle ear function, all ears show (a) measurable EMREOs (mean: 58.7 dB SPL; range 45-67 dB SPL for large contralateral saccades), (b) a phase reversal for contra-versus ipsilaterally-directed saccades, (c) a large peak in the signal occurring soon after saccade onset, (d) an additional large peak time-locked to saccade offset and (e) evidence that saccade duration is encoded in the signal. We interpret the attributes of EMREOs that are most consistent across subjects as the ones that are most likely to play an essential role in their function. The individual differences likely reflect normal variation in individuals’ auditory system anatomy and physiology, much like traditional measures of auditory function such as auditory-evoked OAEs, tympanometry and auditory-evoked potentials. Future work will compare subjects with different types of auditory dysfunction to population data from normal hearing subjects. Overall, these findings provide important context for the widespread observations of visual-and eye-movement related signals found in cortical and subcortical auditory areas of the brain.

## Introduction

Eye movements are critical to linking vision and spatial hearing. Sounds create spatial cues across the head and ears while visual stimuli are detected based on patterns of light impinging on the eye (e.g. e.g. Groh and Sparks, 1992; Blauert, 1997; Groh, 2014). Every eye movement shifts the relative relationship between the visual and auditory reference frames. Thus, information about angle of the eye with respect to head/ears is crucial for integrating the locations of auditory and visual stimuli. However, exactly how this computation occurs is not yet known.

We previously identified a unique type of otoacoustic emission that may play a role in this process (Gruters et al., 2018; Murphy et al., 2020; Lovich et al., 2022). Unlike traditional otoacoustic emissions that can occur spontaneously or in response to auditory stimuli (for review see (Probst et al., 1991; Lonsbury-Martin and Martin, 2003; Shera, 2004), eye movement-related eardrum oscillations (EMREOs) are time-locked to saccade onset and phase-reset at saccade offset (i.e. EMREOs continue after saccades stop with phases that align at saccade offset). Both initial eye position and change in eye position parametrically impact EMREO magnitude and phase (Gruters et al., 2018; Murphy et al., 2020; Lovich et al., 2022), demonstrating the presence of eye-movement-related information at the auditory periphery. This provides the auditory system early access to signals necessary for communication with visual maps of space.

The mechanism(s) underlying such signals may alter auditory transduction in an eye-movement dependent fashion and contribute to previous neurophysiological observations of eye-movement-related modulation of both subcortical (Groh et al., 2001; Zwiers et al., 2004; Porter et al., 2006; Bulkin and Groh, 2012a, b) and cortical (Werner-Reiss et al., 2003; Fu et al., 2004; Maier and Groh, 2010; Willett et al., 2019; O’Connell et al., 2020) stages of the auditory pathway. Indeed, recent work in humans indicates that eye-movement related modulation is widespread throughout auditory-selective regions of cortex (Leszczynski et al., 2023). A full understanding of the eye-movement related signals present at the periphery will ultimately prove vital for understanding these aspects of central processing. .

Our initial study suggested that there are individual differences in the precise nature of this signal (Gruters et al., 2018). Given that all individuals with normal or corrected-to-normal vision, hearing, and eye movements are confronted with the same computational challenge of achieving spatial perceptual stability between vision and hearing across eye movements, the similarities and differences in the EMREOs observed across normal participants can provide clues as to what particular aspects of these signals are relevant to that underlying computational process. Furthermore, like conventional OAEs, EMREOs could eventually have clinical relevance for diagnosing the nature of hearing deficits or auditory-visual integrative dysfunction. Accordingly, in this work, we seek to ascertain which aspects of the EMREO signal are most consistent across normal hearing individuals and which exhibit the most variability. An overarching goal is to establish norms for EMREO data collection and analysis and set the stage for future clinical research in this area.

We report here that certain basic properties of EMREOs were present in all participants: specifically, varying phase and amplitude based on the direction and amplitude of the associated eye movement. Generally, EMREOs are larger when saccades are made to the hemi-field contralateral to the recorded ear than to the ipsilateral field. Individuals varied in the maximum amplitude of their EMREO signals. There was also a slight jitter in the latency of the peaks of the EMREO across individuals, but the pattern of latency changes in the EMREO relating to saccade length/duration is maintained within individual subjects’ responses. Finally, there is greater response variability across subjects for ipsilateral compared to contralateral saccades. Within subjects, EMREOs were highly repeatable across different blocks of trials and recording days.

Identifying and quantifying aspects of EMREO signals that are similar across normal hearing subjects, as we have done here, will ultimately allow comparisons to responses in individuals with different types of auditory system dysfunction. These comparisons should provide insight into the mechanisms that generate and/or modulate the EMREO response as well as provide a means for assessing the feasibility of the EMREO as a future clinical measure of auditory-visual integration.

## Methods

### General overview

This study is part of our ongoing work concerning EMREOs. In our initial study (Gruters et al., 2018) it was unclear if all participants with normal hearing exhibited EMREOs. Our current work involves an ear tip that provides a better seal in the ear canal than the one we used previously (see “Equipment”). With the change in the ear tip, we have successfully recorded EMREOs in all normal hearing individuals who completed an adequate number of trials of a visual saccade task (∼40 subjects). We focus on a subset of these normal hearing individuals for the current study.

Subjects: A total of 10 subjects (20 ears, 6 female, 4 male, 18-39 years old) completed the horizontal-vertical task (Figure 1). All study procedures involving subjects were approved by the Duke Institutional Review Board and all subjects received monetary compensation for their participation in the study. All subjects had hearing thresholds within normal limits (air conduction thresholds < 25 dB HL @250, 500, 1000, 2000 and 4000 Hz, determined through pure tone audiometry using a Maico MI-25 screening audiometer) and normal middle ear function based on standard 226 Hz probe tone tympanometry and acoustic reflexes (Maico touch Tymp MI-24). Screening audiograms, baseline tympanograms, and acoustic reflexes were obtained prior to or on the first day of testing. Middle ear function was assessed with tympanometry on each day of data collection. For all subjects, no significant variation in tympanograms were noted during the testing window.

**Figure 1.**
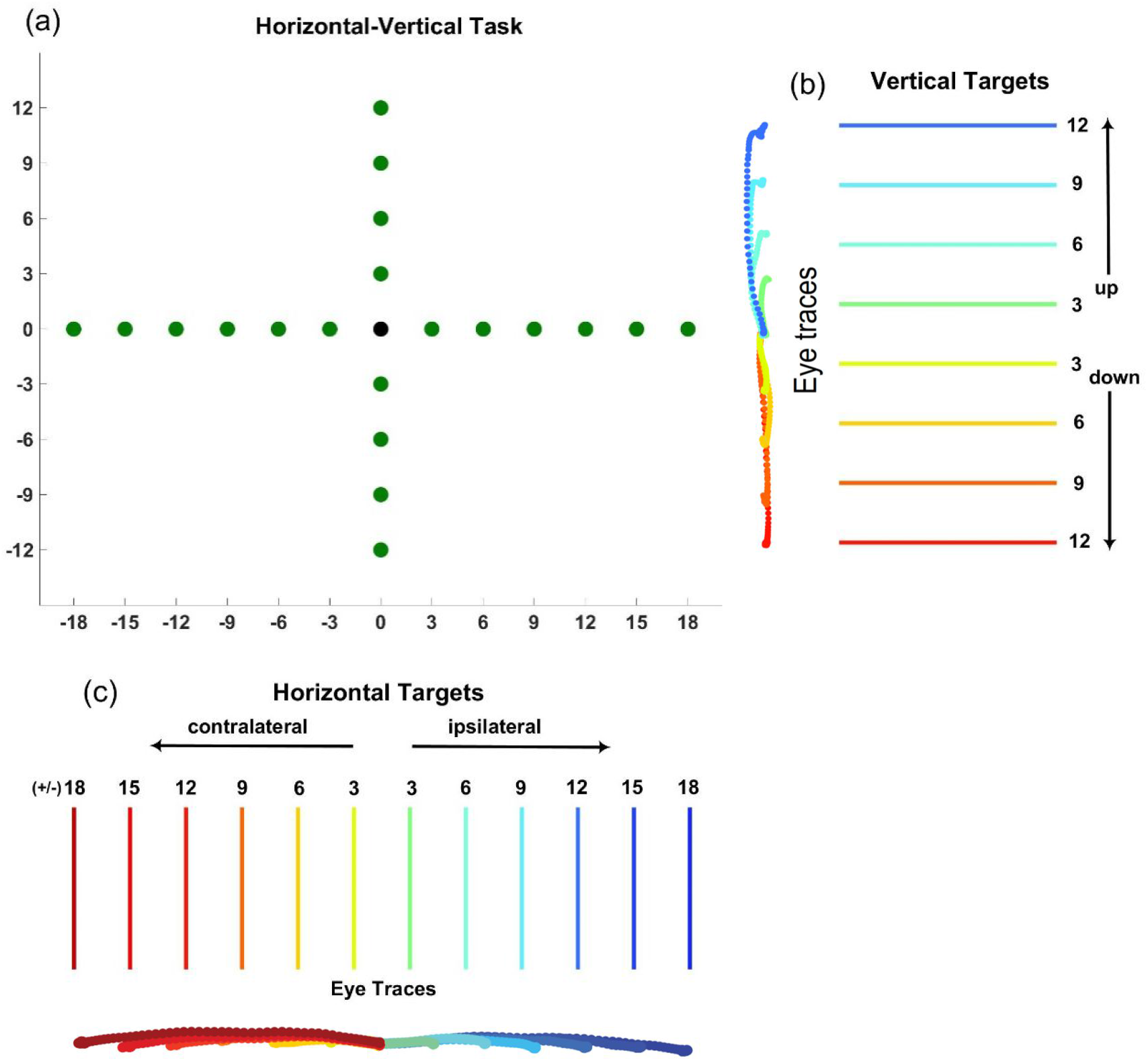
(a) Schematic of horizontal-vertical task and color schemes for subsquent presentation of results as a function of visual target locations. Subjects were asked to fixate on a center point on the screen (black dot) until it disappeared and a target (green dot) appeared. Subjects then saccaded to the target and held fixation until the target turned red, signaling the end of the trial. (b, c) The color code represents saccade direction to target locations. For vertical targets (b), the color code for saccade direction indicates either upward or downward movement and does not change between left and right ear recordings. For horizontal targets (c), the color code for saccade direction is relative to the ears, not to visual space. Contralateral targets are in the hemi-field opposite the ear canal recording and ipsilateral targets are in the same hemi-field as the ear canal recording. Data will be plotted according to this color scheme for *Figures 4 and 5*.

### Tasks

Data are presented here from a visually guided saccade task involving a central fixation point and targets arrayed in the horizontal and vertical dimensions. Targets were spaced 3° apart and spanned a range of +/-18° horizontally and +/-12° vertically (Figure 1). As previously (Lovich et al 2022, Murphy et al 2020), the fixation point was illuminated for 750 ms (until the target appeared). In turn, the green target stayed on for a further 750 ms before turning red, signaling the end of the trial. No external sounds were presented at any time during data collection.

Subjects performed this task while seated in a darkened sound-insulated booth (Acoustic Systems). The stimuli were presented on a monitor positioned 70 cm away; head position was stabilized using a chin rest. Eye position and ear canal microphone data were recorded simultaneously (see Equipment) while subjects performed the saccade task. Data collection sometimes occurred on multiple days, and involved 1-3 one-hour sessions per subject. Recordings were made in blocks of ∼125 trials each, presented pseudorandomly. Each test session consisted of 3-6 blocks, with breaks offered between blocks. Recording time for each block was approximately five minutes and each test session lasted approximately one hour, including instructions, eye calibrations, microphone assessments, and rest periods. Data were combined across blocks and sessions. See Table 1 for additional details regarding sessions, blocks, and included trials per subject.

**Table 1.** Summary of amount of data per subject.

| *Subjects* | *Number of days* | *Number of blocks* | *Number of trials* | *N trials included after saccade screens* | *N trials included after saccade and microphone noise screens* | *N trials included per target (20 targets)* | *Overall % included* | *Overall % excluded due to microphone noise screens* |
| --- | --- | --- | --- | --- | --- | --- | --- | --- |
| S05 | 1 | 4 | 488 | 448 | 438 | 21.9 | 89.8% | 2.0% |
| S15 | 2 | 11 | 1094 | 832 | 813 | 40.7 | 74.3% | 1.7% |
| S17 | 1 | 3 | 326 | 287 | 285 | 14.3 | 87.4% | 0.6% |
| S24 | 1 | 5 | 509 | 432 | 430 | 21.5 | 84.5% | 0.4% |
| S45 | 2 | 9 | 550 | 425 | 409 | 20.5 | 74.4% | 2.9% |
| S49 | 2 | 10 | 758 | 574 | 556 | 27.8 | 73.4% | 2.4% |
| S50 | 1 | 4 | 447 | 341 | 339 | 17.0 | 75.8% | 0.4% |
| S52 | 1 | 4 | 439 | 405 | 400 | 20.0 | 91.1% | 1.1% |
| S71 | 2 | 8 | 747 | 637 | 635 | 31.8 | 85.0% | 0.3% |
| S73 | 3 | 10 | 986 | 749 | 746 | 37.3 | 75.7% | 0.3% |
| **Average across subjects** | **1.6** | **6.8** | **634.4** | **513** | **505.1** | **25.3** | **81.1%** | **1.2%** |

### Data collection equipment

The entire data collection process was managed through MATLAB 2017b (MathWorks) and Psychtoolbox on a Mac Pro computer.

Microphone recordings were obtained in both ear canals simultaneously during all testing sessions using Etymotic Research ER10B+ Low Noise Microphone Systems and ER10-14 foam ear tips, coupled with ER-2 insert earphones. A Focusrite Scarlett 2i2 low-latency audio interface provided audio capture and playback at 48 kHz sampling rate through the Etymotic system. Microphone data were downsampled from 48 kHz to 2 kHz prior to analysis to reduce processing time using MATLAB’s “resample” function (MathWorks). This is well above the Nyquist frequency for the range of interest (<100 Hz). Except where noted, each subject’s microphone recording values are Z-scored or expressed in units of standard deviation relative to the variability observed in a 30 ms baseline window starting 150 ms prior to saccade onset or offset, i.e. during a period of steady fixation. The individual microphone samples during this baseline were pooled across all the trials within a recording block and their mean and standard deviation were used to compute the Z-scored values for each of the individual trials in that same block.

To ensure stable microphone function across all testing sessions, the Etymotic ER-2 earphones were used to play a sequence of tones prior to each test session as well as before each test block (pure tone sweeps from 10-1600 Hz: 10 Hz steps for 10-200 Hz, 100 Hz steps for 300:1000 Hz, 200 Hz steps for 1200-1600 Hz). Before the subject arrived, we assessed the function of the microphone system using a device to simulate average ear canal volume (A 3 cc syringe tip was cut off and replaced with an ear tip typically used with the microphone system. The syringe stopper was pulled to a volume of 1.25 cc and approximately 0.25 cc of cotton wadding was added to simulate the soft tissue of the ear canal). Microphone measurements were considered valid if the input-output of the tone sequence during this pre-testing evaluation routine matched the standard set obtained during pre-testing evaluations.

The tone sequence also played prior to each test block, with microphones placed in the subject’s ears, to evaluate microphone response stability throughout the testing session. Microphone output recorded during tone sweeps was monitored. Any changes (e.g. an earbud dislodging) caused the associated recording block to be discarded.

Eye tracking information was recorded with the SR Research Eyelink 1000 system at a 1 kHz sampling rate. Eye tracking was calibrated at the beginning of each test session using the Eyelink 1000 calibration program and again if the subject moved position or paused for a break in data collection.

### Analysis

As in earlier work (Lovich et al., 2022; Murphy et al., 2020), we smoothed the eye tracking data using a lowpass discrete filter with a 7 ms window. Microphone recordings were synchronized with eye tracking data at saccade onset and offset, defined based on the third derivative of eye position (jerk). The point where change in eye acceleration was greatest was marked as saccade onset while the point where change in eye deceleration was greatest was marked as saccade offset. Trials were first excluded based on saccade performance and then on quality of the microphone recordings. For eye tracking, trials were excluded if subjects made more than a single saccade to reach the target, if eye tracking signal was lost during the trials (e.g., from blinks), if eye movement was significantly slower than an average saccade (specifically, the maximum velocity was less than 50 degrees/second), or if the saccade curved more than 4.5°. Finally, a minimum fixation of 200 ms before and after the saccade was required in order to include a trial in analysis. The majority of trials that did not meet criteria for inclusion were due to issues with eye tracking, eliminating 19% of trials. This number is slightly higher than previously reported from our lab, likely due to the specifics of this task (visual targets for the horizontal-vertical task are more closely spaced, with only 3° separation, compared to earlier tasks with 6° target separation). Once the criteria for eye tracking were met, a modest number of trials (1%) were excluded based on integrity of microphone recordings. Individual trials were excluded when the whole trial RMS value was >10 SD than the average RMS value for the entire trial block. Following application of exclusion criteria, there were between ∼14-41 trials per target for each subject (average of 25.3 trials per target for each subject, see Supplementary Table 1 for details). The Z-scoring of the microphone values described above under “Data collection equipment” was computed after these exclusion criteria were applied.

### Regression analysis

We have used a general linear regression approach similar to those used in functional magnetic resonance imaging (fMRI) research (e.g., Monti, 2011) in our related studies (Gruters et al., 2018; Lovich et al., 2022; Lovich et al., 2023) and we take a similar approach here. We fit the microphone data for each individual subject’s ear using the following equation:

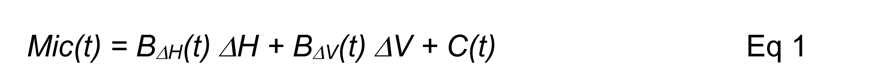

The variables *ΔH* and *ΔV* correspond to the respective changes in eye position associated with a given trial (one value of each per trial) and were measured in degrees. The coefficients *B_ΔH_*and *B_ΔV_*vary with time in the trial^1^ and reflect the dependence of the microphone signal on the respective eye movement parameters. The Z-scoring described above causes these coefficients to have units of standard deviation per degree. The term *C(t)* contributes a time-varying “constant” independent of eye movement metrics, with units of standard deviation. lt can be thought of as the best fitting average oscillation across all eye movements. We used the measured values of change in eye position for this analysis rather than the associated fixation and target locations so as to incorporate trial-by-trial variability in fixation and saccade accuracy for a given visual target (Lovich et al., 2022).

We determined whether an individual subject’s ear exhibited a statistically significant EMREO in a two-stage process. First, we evaluated the time-varying p-values of the regression fits p(t) during a window starting 5 ms before saccade onset and ending 70 ms after saccade onset; this corresponds to 149 data points given the downsampling of the microphone data. These data points are not independent of one another, so conventional Bonferroni correction would not be appropriate. Accordingly, we evaluated whether the

The key strengths of this regression approach are its relative simplicity given the many unknowns regarding the EMREO phenomenon and its flexibility for use across studies that use different task designs/target locations. The main weaknesses are that (a) the relationship between microphone voltage and eye movement parameters may not be linear; (b) the microphone signals may not be normally distributed; (c) the oscillatory nature of the signal means that zero crossings occur and thus the fitted coefficients will not differ from zero during those periods of time; (d) as noted above, data points are correlated across time so correcting for multiple comparisons is not straightforward. However, points (a-c) all serve to reduce the likelihood of observing “significant” p values; and point (d) is not expected to introduce a bias in favor of significance vs. non-significance. If results are borderline and interpretation is uncertain, results can be compared to a shuffled control, which we have done previously (Gruters et al., 2018).

### Estimating EMREO sound amplitude

The true sound pressure levels produced during the EMREO are difficult to ascertain because the EMREO’s frequency of ∼40 Hz is well outside the frequency range for which most standard audio components are calibrated (typical lower limit of about 100-200 Hz). We have addressed this problem previously by estimating the EMREO amplitude based on a model of the low frequency-response characteristics of the ER10B+ microphone (Christensen et al., 2015; Gruters et al., 2018). Here, we took a different approach. Using a Bruel & Kjaer sound level meter (Model 2245, reported by the manufacturer to have a flat frequency response from 20 Hz to 20 kHz, Z-weighting), we measured the sound pressure levels associated with the 40 Hz pure tones generated by the Etymotic ER-2 transducers as described above. These measurements were made in a short length of tygon tubing simulating the approximate volume and sound-absorption properties of the human ear, and were simultaneously recorded with the ER10B+ microphones. This provided a voltage-to-sound-pressure-level mapping in the 40 Hz frequency range. We then used these values to estimate the sound pressure level of the EMREO signal in individual subjects (see Figure 6C).

## Results

### Statistical significance of individual EMREOs via regression

Regression is a powerful method for evaluating the overall pattern of dependence of the EMREO waveform on saccade parameters, and provides for an estimate of statistical significance at the individual subject level. All subjects tested in this study showed statistically significant EMREOs per the test described in *Methods: Regression analysis*. Briefly, this involved assessing the influence of the horizontal and vertical displacement of the eyes on the recorded microphone signal via time-varying regression (Eq 1). We evaluated the time varying p-values of this regression (p(t)) during a window of -5 to 70 ms, when aligned to saccade onset. Any ear that exhibited p(t)<0.05 for more than 10% of this time window was considered statistically significant. As shown in Table 2, all 20 ears in 10 subjects exhibited statistically significant EMREOs when aligned to saccade onset (range 21-85%). A similar process was adopted to evaluate saccade offset-aligned EMREOs, with an evaluation window of -25 to 50 ms before/after saccade offset. The range of observed values was 22-83%, again reaching significance for all 20 ears. (Table 2).

**Table 2.** Percentage of analysis time window where p-values of the regression fits were less than 0.05. Significance for greater than 10% of the analysis window was used to determine the presence of an EMREO for that ear (onset-aligned analysis window: -5 to 70 ms relative to saccade onset; offset-aligned analysis window: -25 to 50 ms relative to saccade offset; see Methods).

| Onset-aligned | Offset-aligned |
| --- | --- |

*Table 2. Percentage of analysis time window where p-values of the regression fits were less than 0.05.
|  | Left Ear | Right Ear | Left Ear | Right Ear |
| --- | --- | --- | --- | --- |
| S05 | 84.6% | 42.2% | 83.2% | 49.0% |
| S15 | 54.4% | 45.0% | 28.2% | 33.6% |
| S17 | 53.7% | 26.1% | 69.1% | 24.2% |
| S24 | 51.7% | 21.7% | 47.0% | 51.7% |
| S45 | 57.0% | 40.2% | 40.9% | 38.3% |
| S49 | 52.3% | 26.9% | 26.8% | 40.9% |
| S50 | 63.8% | 62.2% | 55.0% | 37.6% |
| S52 | 64.4% | 61.0% | 59.1% | 68.5% |
| S71 | 81.2% | 42.6% | 49.0% | 21.5% |
| S73 | 23.5% | 38.2% | 22.1% | 33.6% |

### Horizontal and vertical coefficients of regression fits across time

Each coefficient of the regression equation can be plotted over time. Figure 2 shows the slope coefficients for horizontal eye displacement (left column), vertical eye displacement (middle column) and the time-varying constant (right column), aligned to saccade onset. In the left column, all subjects show an oscillatory pattern in relation to the horizontal displacement of the eyes with the peaks of the oscillations significantly different from zero for all subjects (thick lines, p<0.05). Note the phase of the oscillations is opposite for the two ears, thus encoding horizontal saccade direction relative to the recorded ear.

**Figure 2.**
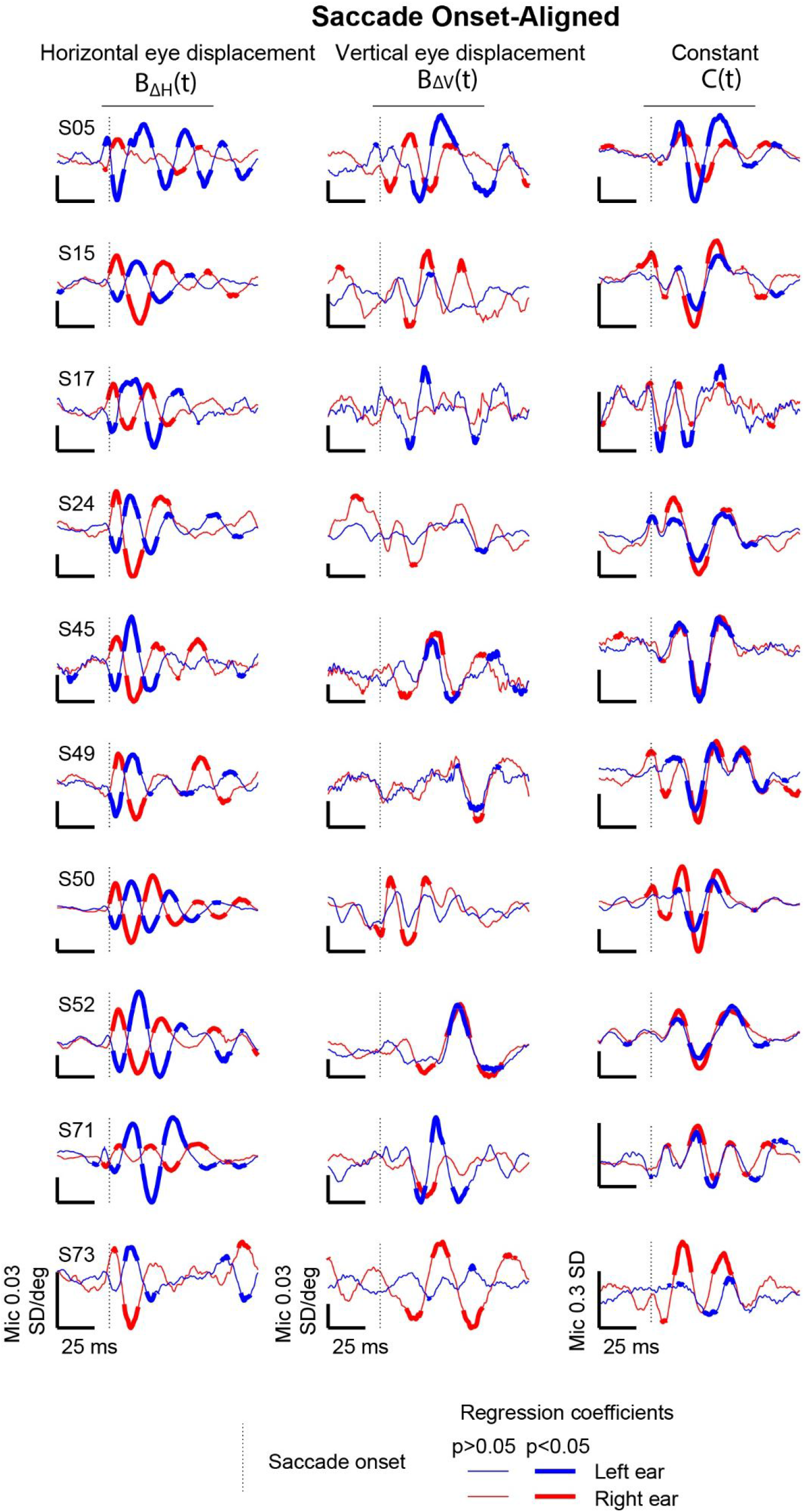
Coefficients of linear regression fits for individual subjects, aligned to saccade onset. Blue traces are left ear data and red traces are right ear data. Thick line segments indicate epochs of time in which the corresponding regression coefficient differed from zero (p<0.05). The time window for analysis of aggregate statistical significance as provided in Table 1 is marked by the horizontal black bar at the top of each column (-5 to 70 ms with respect to saccade onset). Y-axis units are standard deviations per degree (horizontal and vertical eye displacement columns) or simply standard deviations (“constant” column) and vary across panels as indicated by the calibration bars. See Supplementary Figure 1 for individual subject data plotted on the same graphs for comparison purposes.

Compared to the horizontal component, the time where vertical eye displacement (middle column) differs significantly from zero is much shorter and the latencies of the significant regions are not the same across subjects. However, all subjects have a waveform peak within the EMREO analysis time window that is significantly different from zero for at least one of the two ears.

For the time-varying constant component (right column), several subjects show tri-phasic waveforms (S05, S45, S52), while the remaining subjects show multiple positive and negative peaks that are significantly different from zero in one or both ears. This portion of the regression fits can be thought of as the basic EMREO waveform upon which dependency on horizontal and vertical eye displacement is superimposed.

Figure 3 shows the same data as Figure 2, but aligned to saccade offset. As before, the oscillatory pattern in the horizontal displacement component (left column) is seen following saccade offset and the peaks of the oscillations are significantly different from zero (thick lines, p<0.05) for all subjects and all ears. The vertical eye displacement component (middle column) reveals even greater variability across subjects compared to corresponding vertical onset-aligned responses. The time-varying constant (right column) is also less predictable across subjects when aligned to saccade offset than when aligned to saccade onset.

**Figure 3:**
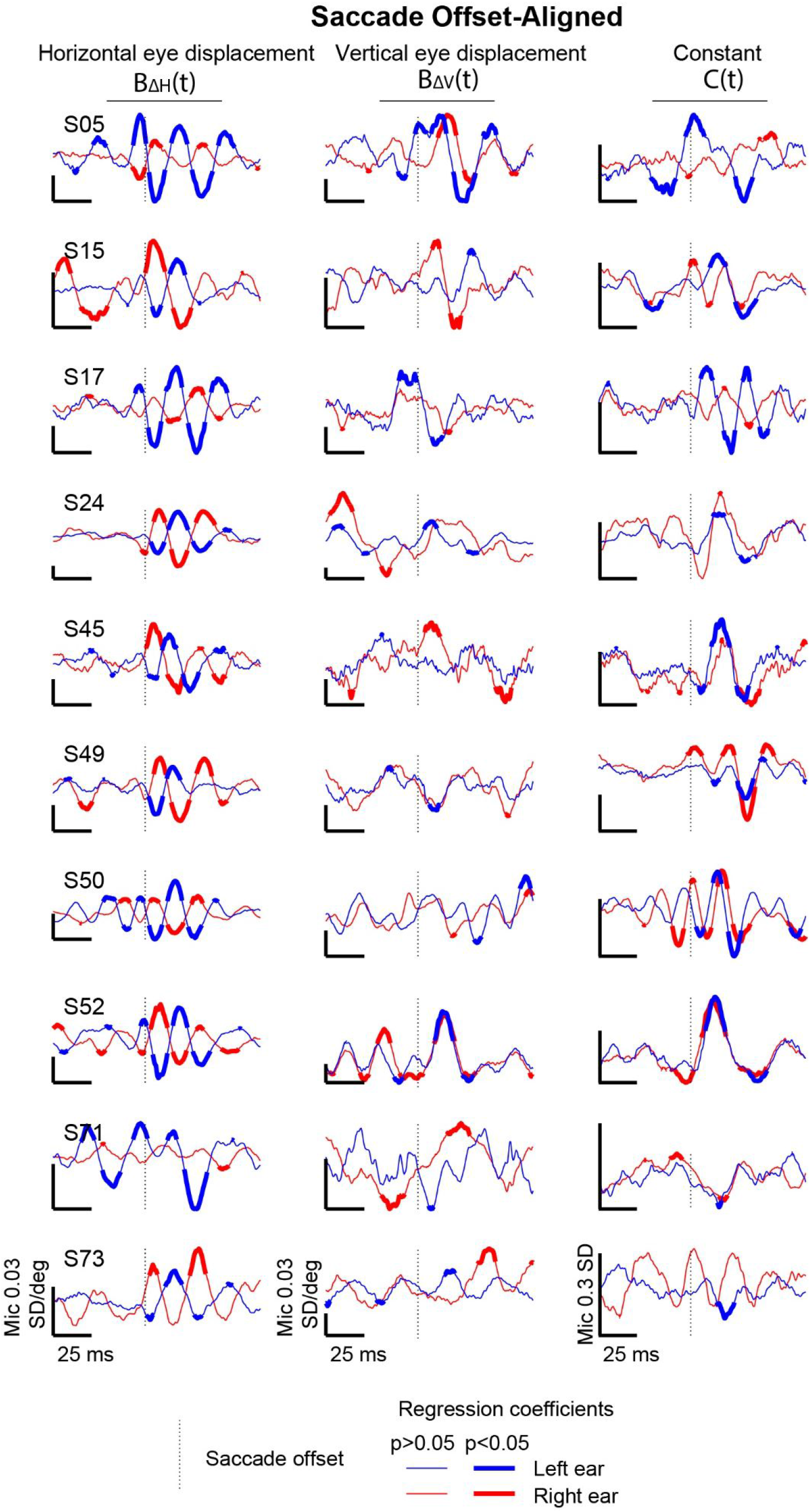
Coefficients of linear regression fits to saccade-offset-aligned responses. Conventions same as Figure 2. The time window for evaluation of aggregate statistical significance of regression coefficients (Table 1) was defined as -25 to 50 ms with respect to saccade offset as denoted by a black bar at the top of each column. Thick lines indicate when the waveform differs significantly from zero (p<0.05). See Supplementary Figure 1 for individual subject data plotted on the same graphs for comparison purposes.

Overall, for both onset-and offset-aligned responses, the duration of statistical significance is generally greater in relation to horizontal eye displacement compared to the other two regression coefficients. In other words, the confidence intervals for the coefficient concerning horizontal eye displacement differ from zero for a greater fraction of the time during the evaluated time windows (-5 to 70 ms with respect to saccade onset and -25 to 50 ms with respect to saccade offset) in relation to vertical displacement and the time-varying constant.

### EMREO waveforms by target location

A deeper dive into the EMREO waveforms for each target location confirms the general patterns observed via regression. Since EMREOs are detected in all ears tested, and since the temporal patterns are largely similar across ears within individual subjects, we combined right and left ear microphone data as a function of target location with respect to the recorded ear. Therefore, rightward/leftward targets for left ear recordings were combined with corresponding leftward/rightward targets for right ear recordings, maintaining the contralateral/ipsilateral distinctions.

Figure 4 shows each subject’s microphone responses to both horizontal (1^st^ column) and vertical (2^nd^ column) saccade targets, aligned to saccade onset, and color-coded according to the scheme illustrated in Figure 1. As seen with the regression fits, all subjects show oscillations that invert phase with change in horizontal saccade direction (red-orange-yellow traces are from saccades to contralateral targets; indigo-blue-green traces are to ipsilateral targets). For all horizontal target locations, oscillations begin close to saccade onset and have a similar waveform pattern both within and across subjects for approximately the first 30 ms. There is a small peak/trough at ∼5 ms after onset, followed by larger peaks/troughs at ∼15 and ∼30 ms. The amplitude, latency and phase direction of the waveform peaks are synchronized across all subjects. Although the oscillations continue beyond 30 ms, the pattern becomes less predictable across subjects: the consistent phase reversals for contralateral/ipsilateral targets do not persist for most subjects, and waveform amplitude is reduced until returning to pre-saccade baseline levels around 100 ms after saccade onset. (The waveform variations by target location observed in the time window between 40 to 90 ms after saccade onset will be discussed later, in Figure 8).

**Figure 4.**
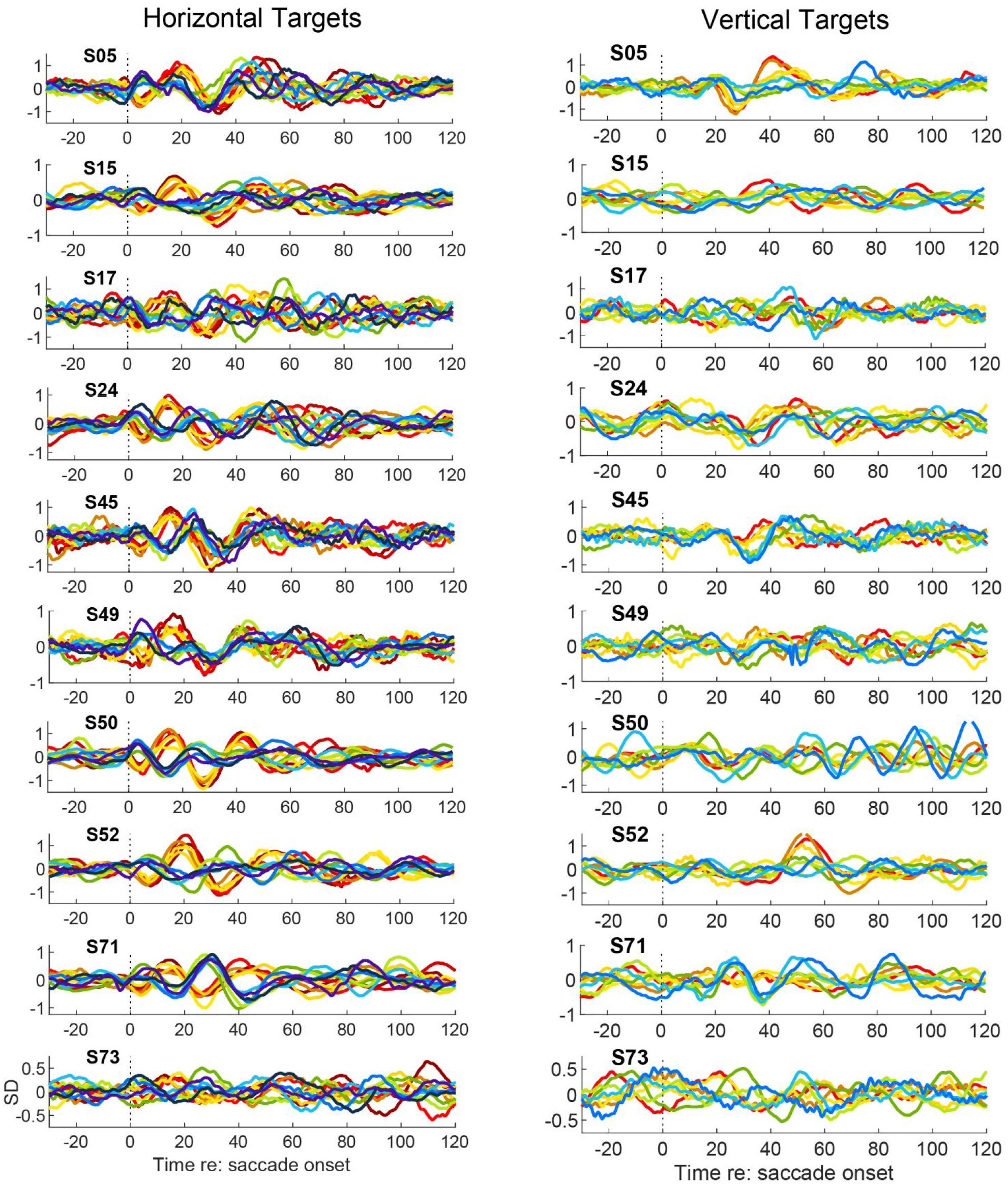
EMREOs recorded during saccades to horizontal targets show more consistent patterns within and across subjects compared to responses during saccades to vertical targets. Average microphone waveforms, z-scored and aligned to saccade onset, are shown for all subjects (y-axis units = standard deviation relative to baseline variation). Each waveform trace is a subject’s averaged microphone response to a single target (both ears combined; see *Figure 1* for color code). Red-orange-yellow traces are responses during saccades to contralateral targets and indigo-blue-green traces are to ipsilateral targets. For horizontal targets, in the first 30 ms after saccade onset, every subject has the expected phase inversion with change in saccade direction and amplitude; latency and phase direction of peaks are aligned. In the 40 to 90 ms time window, which coincides with saccade offsets, there are peaks with latency shifts that appear to align to saccade duration. For vertical targets, no consistent pattern in waveform amplitude or latency can be discerned across subjects.

As observed with the regression fits, the oscillation patterns for purely vertical saccades are much less predictable across subjects. The latency of the largest peak in the oscillation to vertical saccades is variable across subjects. There is no consistent pattern of phase inversion with changes in saccade direction (up vs. down). In addition, latencies of the maximum peak responses do not show a predictable pattern across subjects. Overall, there is greater variability in response latencies and amplitudes during vertical saccades compared to responses during horizontal saccades.

The EMREO waveforms following saccade offset show similar properties as seen in the saccade onset-aligned data (Figure 5). As with onset-aligned responses, EMREOs are especially evident for saccades in the horizontal dimension (left column). A phase-reversed pattern for contra-vs. ipsi-versive saccades is evident in all subjects, although the time course and latency of the peaks varied across subjects. EMREOs lasted for at least 40 ms after horizontal saccade offset in all subjects, and are present to approximately 60 ms after saccade offset in several subjects (e.g. S05, S17, and S73). As with saccade onset-aligned data, inter-subject variability in saccade offset-aligned responses is greater for vertically directed saccades than for horizontally directed saccades. Generally, the waveforms appear less stereotyped across vertical target locations both within and across subjects, just as they do for saccade onset-aligned waveforms.

**Figure 5:**
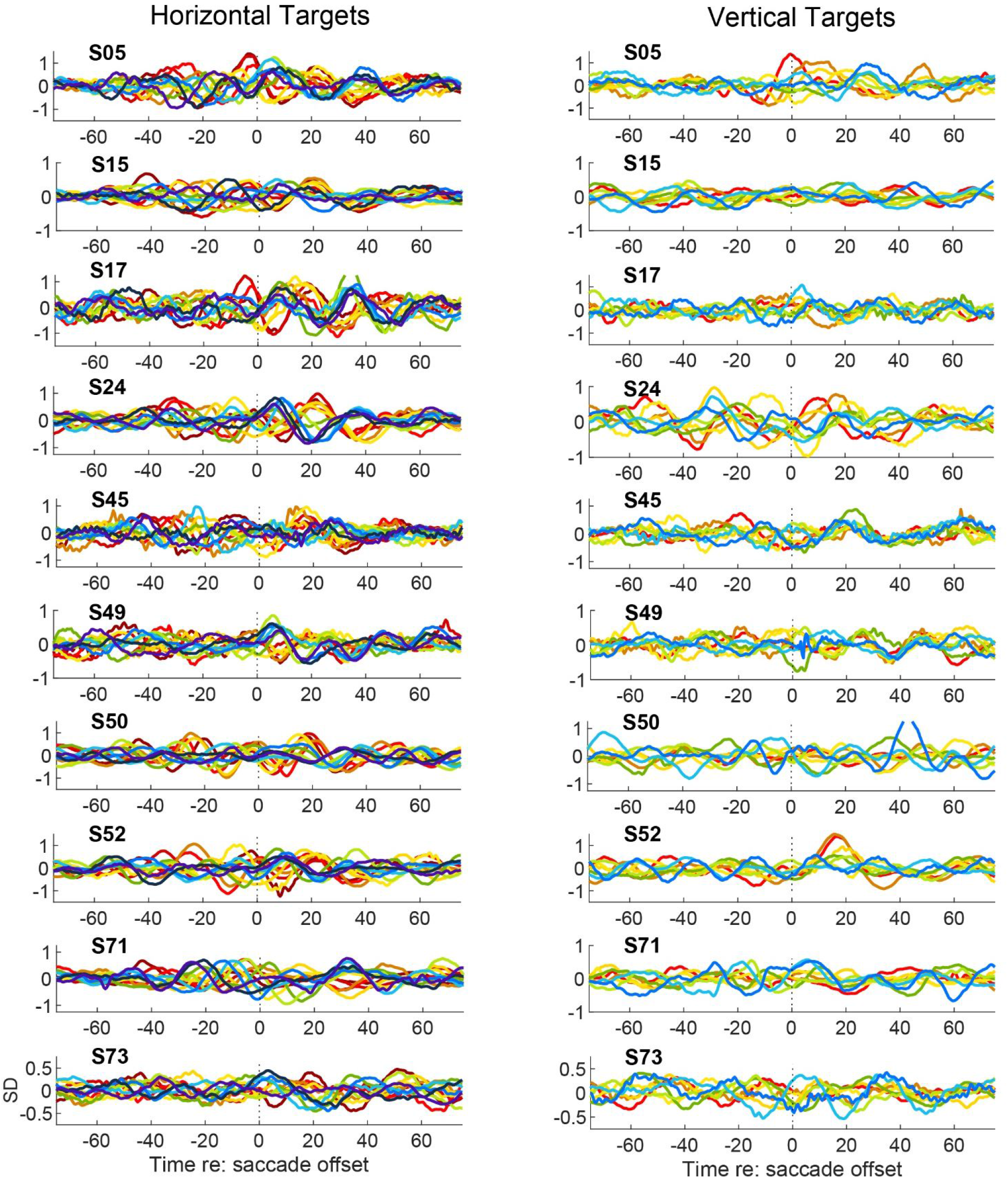
Similar to *Figure 4* but aligned to saccade offset. Offset responses to horizontal saccade targets reveal patterns similar to those seen in onset responses but are not as well organized. A phase inversion of the response with change in saccade direction is seen in all subjects. The waveforms align with the first peak/trough around 5 ms and the second peak trough around 20 ms after saccade offset. The responses return to pre-saccade baseline levels by approximately 60 ms in all subjects. Vertical offset responses are quite variable across subjects and similarities in response patterns across subjects do not easily emerge.

### EMREOs for contralaterally-directed saccades are larger and less variable than ipsilateral ones

Although EMREOs that occur during horizontal saccades are less variable than those during vertical saccades, the EMREOs associated with saccades within the horizontal dimension were not fully symmetric across the contralateral vs. ipsilateral directions. Two types of differences were found. First, the amplitude of the EMREO waveform was slightly larger for contralaterally-directed vs. ipsilaterally-directed saccades. This can be seen in Figures 4 and 5, where the contralaterally-directed (red-orange-yellow) traces exhibit larger oscillations than the ipsilaterally-directed (indigo-blue-green) traces for individual subjects. Second, there was greater variability in the EMREOs associated with the ipsilateral targets. This variability is evident in both amplitude and latency. Not only were the EMREOs associated with saccades to ipsilateral targets smaller on average than those to contralateral targets, but they also exhibited greater inter-subject variation in EMREO amplitude, whereas the EMREOs associated with contralateral saccades were quite consistent across subjects. Furthermore, the latencies of the peaks/troughs are more precisely aligned across subjects’ responses to contralateral targets than is the case for ipsilateral targets.

The contralateral vs. ipsilateral amplitude differences and the amplitude/latency variation are illustrated and quantified further in Figure 6. Figure 6A-B shows the average microphone signals for individual subjects for the 18 degree horizontal contralateral (A) vs. ipsilateral (B) target locations aligned on saccade onset. The contralateral waveforms are larger and less variable across subjects than the ipsilateral ones. Converting the underlying raw data to units of peak-equivalent sound pressure level (see Methods), the mean sound level for EMREOs to 18 degree contralateral targets is 58.7 dB SPL, whereas for ipsilateral targets it is 55.7 dB SPL (Figure 6C); this 3 dB difference corresponds to the approximately 30% difference in amplitude visible in panels A vs B where the data are expressed in Z-score units (standard deviation). The range of amplitudes is about 45-67 dB SPL across subjects.

**Figure 6.**
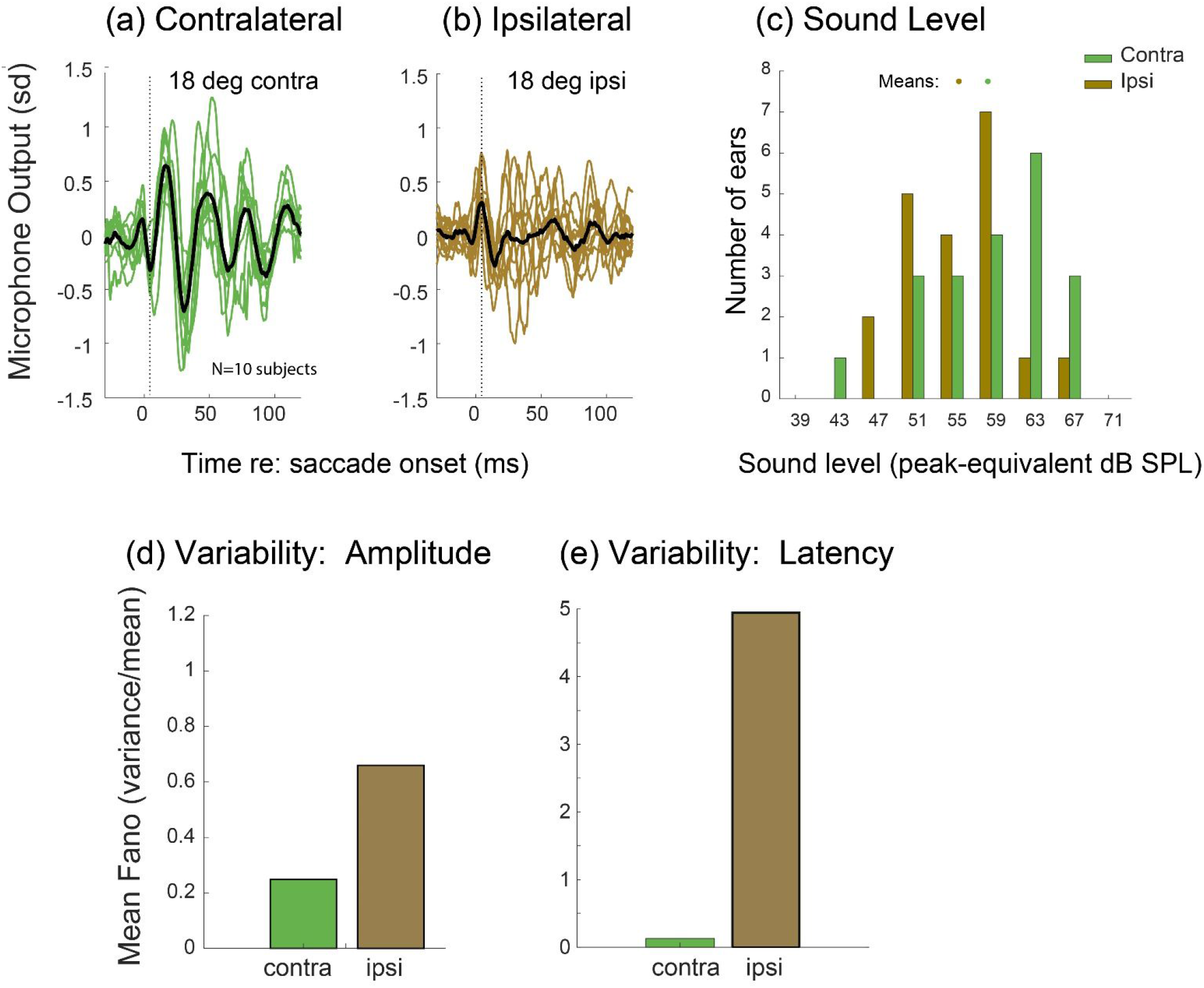
Peak/trough amplitudes and latencies are more consistent across subjects during saccades to contralateral than ipsilateral targets. (a-b). Each trace is an individual subject’s averaged microphone response to the 18 degree contralateral (a) or ipsilateral (b) horizontal target location, aligned to saccade onset. Waveforms are more similar across subjects for contralateral targets than for ipsilateral targets. Overall, amplitudes of ipsilateral responses appear slightly smaller in individual subjects compared to contralateral responses. (c). Sound level of EMREOs associated with the 18 degree contralateral or ipsilateral target locations for individual subjects (left and right ears computed separately; see “Methods: Estimating EMREO sound amplitude” for details. (d-e) The variance-to-mean ratio of EMREO amplitude (d) and latency (e) is smaller for contralateral vs ipsilateral horizontal target locations, where amplitude and latency were computed based on the maximum peak (contralateral) or trough (ipsilateral) observed in a window 0-35 ms after saccade onset (the peaks and troughs used for these analyses were expressed in units of Z-scores not dB). Both amplitudes and latencies of ipsilateral responses are more variable across subjects compared to contralateral responses

The greater cross-subject variability of both peak amplitude and timing for ipsilateral vs contralateral saccades is further quantified in Figure 6D-E. For both peak amplitude (D) and latency (E), the cross-subject variance-to-mean ratio is higher for ipsilateral (brown) than for contralateral (green) saccades. While we note here that the overall amplitude of the ipsilateral responses are smaller than contralateral, the greater temporal variability across subjects for ipsilateral saccades is an additional underlying factor determining what appears to be *substantially* smaller EMREO amplitudes to ipsilateral visual targets reported in group-averaged population data (Lovich et al 2022).

### Relationship between EMREOs and horizontal saccade length/duration

The alignment of EMREOs on both saccade onset and saccade offset implies a relationship to saccade duration. Saccade duration is known to co-vary with amplitude: to move the eyes to a new position as fast as possible, the brain causes the extraocular muscles to initially apply nearly the same amount of force per unit time for all saccades regardless of their amplitude, and simply stops the movement sooner for shorter vs. longer amplitude saccades (roughly speaking). Does the EMREO show a corresponding pattern? To investigate this question, we focused on the EMREOs to contralateral targets given that they exhibited the least variable amplitudes and latencies across subjects (e.g. Figure 6D). Figure 7A shows the grand-average microphone responses (mean waveform of all subjects’ averaged waveforms) to each contralateral horizontal target, aligned to saccade onset (colored dots indicate the timing of saccade offsets). The latencies of the waveform peaks and troughs align across all target locations during the first 35 ms after saccade onset (which is during the saccade for all but the shortest duration saccades). Peak EMREO amplitudes increase with saccade amplitude during this period, but the effect is relatively modest. After this initially temporally-aligned peak, there is a systematic delay in the latencies of subsequent peaks with increasing saccade amplitude (i.e., in the 40 to 90 ms time window following saccade onset). These findings show saccade amplitude is encoded in two ways: first, via the amplitudes of the initial short latency peaks/troughs within 30 ms after saccade onset; second, via a series of temporally delayed peaks/troughs seemingly related to saccade duration. Overall, this combination of relatively modest differences in initial amplitude, paired with substantial differences in latency and duration, are strongly reminiscent of the known temporal/amplitude patterns of force generation of oculomotor saccade commands (Bahill et al., 1975; Baloh et al., 1975; Van Opstal and Van Gisbergen, 1987).

**Figure 7.**
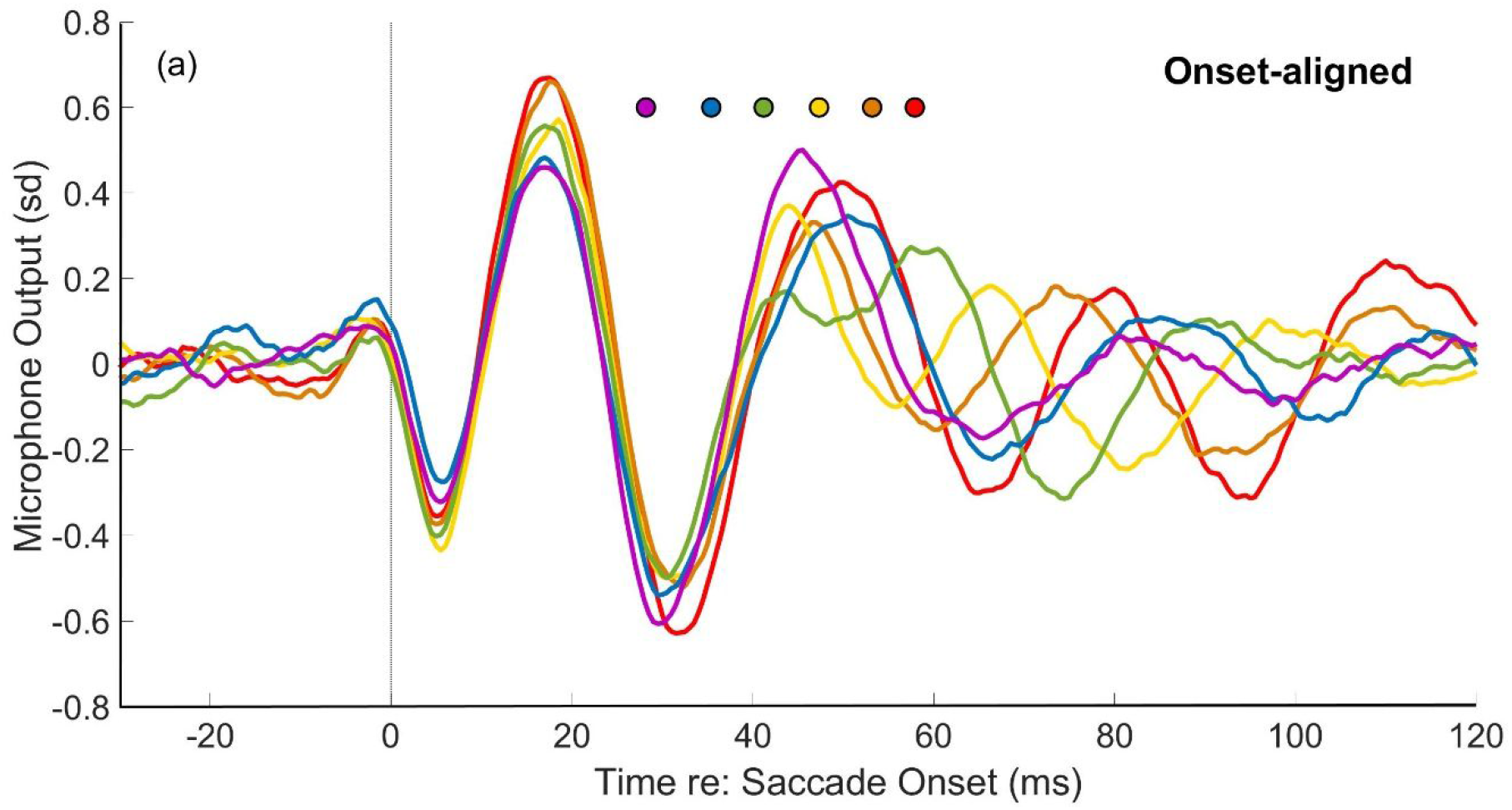
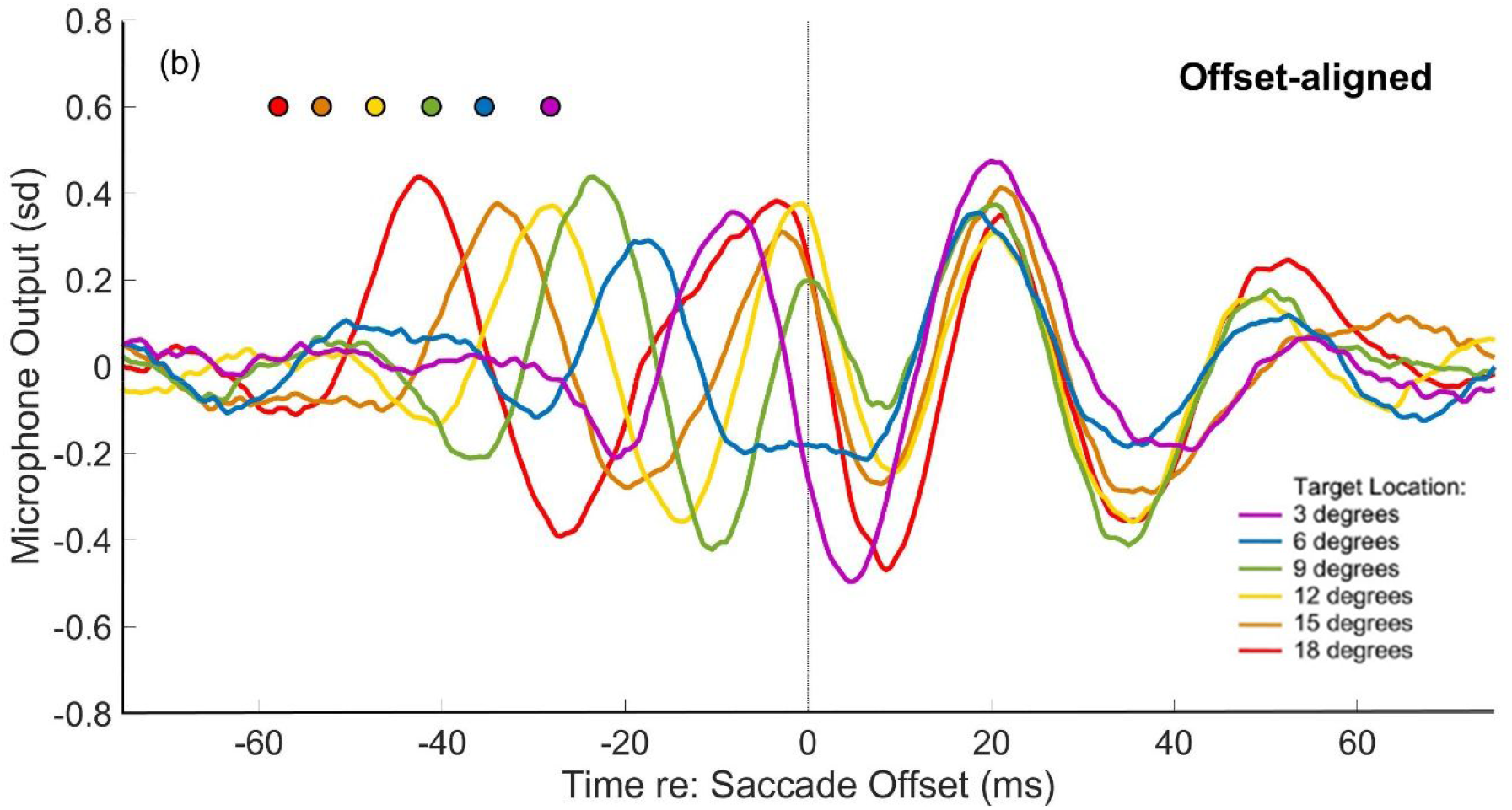
Relationship between timing of EMREO peaks and saccade onset/offset. (a) Grand-averaged waveforms (n=10) to contralateral horizontal visual targets aligned to saccade onset. The latency of the first peak (i.e. in the 0-to-40 ms time window) is consistent across different target locations (different saccade durations), but later peaks (i.e. 40 to 90 ms after onset) are delayed by amounts that depend on target location (different colors) and corresponding saccade duration (colored circles above the waveforms). (b) Same data as in panel (a) but aligned on saccade offset. The first positive-going peak after offset occurs at approximately 20 ms for all target locations.

This pattern is confirmed when viewing the same grand-average waveforms aligned on saccade offset (Figure 7B; colored dots indicate the timing of saccade onsets). In the 50 ms time window prior to saccade offset, there is a systematic shift in the latencies of the peaks/troughs that roughly corresponds with saccade onsets for each target location. Following saccade offset, the peak latencies realign. However, the systematic increase in peak amplitude with increase in saccade magnitude that was seen during the saccade is less evident during this period and the waveform returns to baseline levels by 60 ms after saccade offset.

### Individual variation in EMREO peak latencies and saccade length/duration

Might the variation in EMREOs across subjects be related to individual variation in saccade duration? While saccades are highly stereotyped eye movements, exhibiting characteristic velocity/duration/amplitude relationships within individuals, there can be variation across individuals. To consider this question, using onset aligned data, we identified the first positive EMREO waveform peak after saccade offset for each subject and target, averaged across trials. As shown in Figure 8a, there is a linear relationship between the latency of this peak relative to saccade onset (y-axis) and saccade duration (x-axis). The overall pattern is consistent with the pattern illustrated via population grand-average in Figure 7, but here we can also see that the data points for individual subjects cluster along the same regression line for the whole population. Note the clustering by color, indicating that saccade duration is determined by target location. Different subjects’ data points are shown via different symbols. Although there is overlap in saccade duration for adjacent targets, the pattern for each subject is rather consistent. For example, the subject denoted with an equilateral triangle has relatively short saccade durations for each target and has the shortest peak latency for each target location. Similarly, the subject denoted with a square has relatively long saccade durations and the longest latencies for target locations.

**Figure 8.**
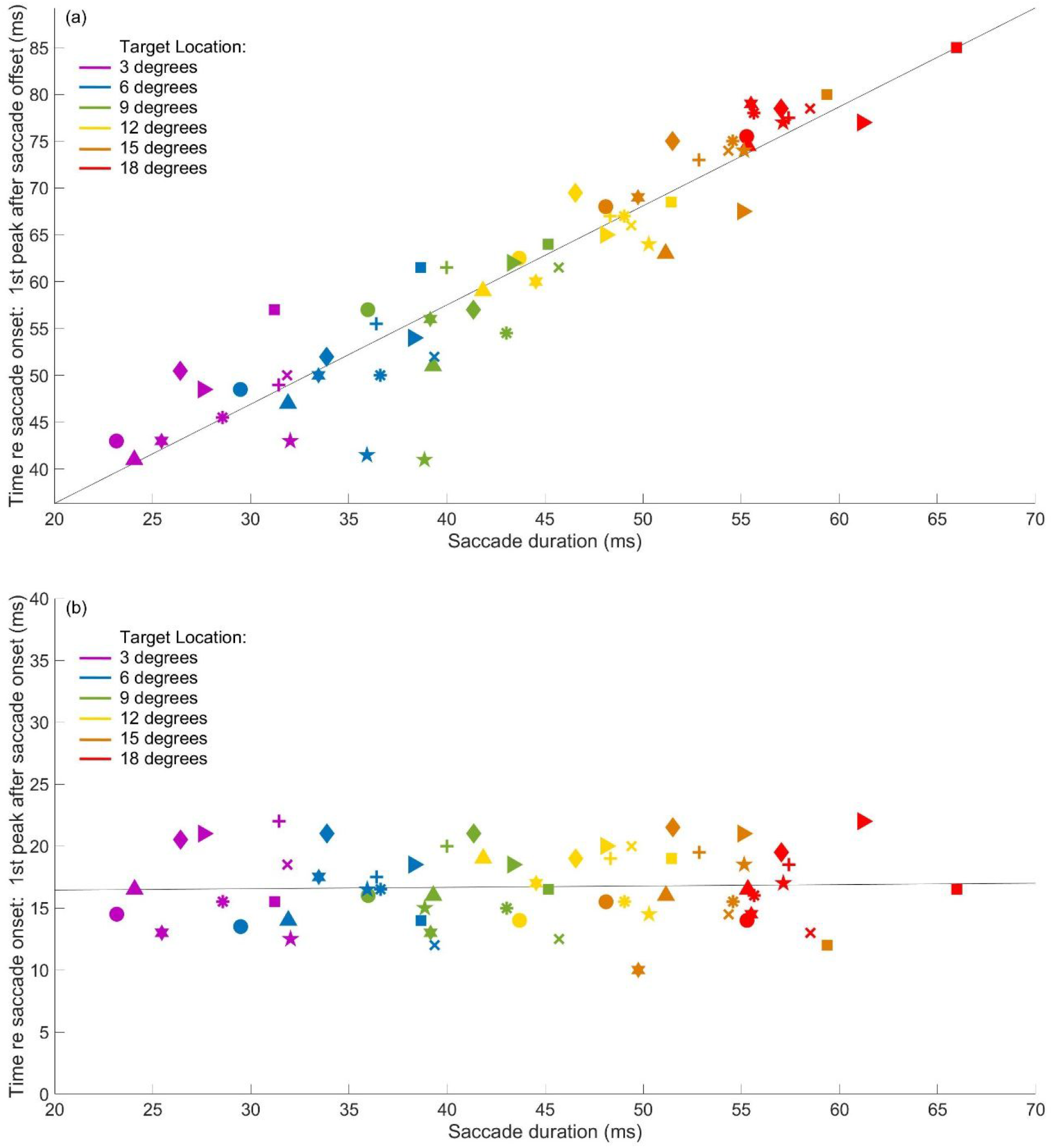
(a) Individual variation in saccade duration is associated with individual variation in EMREO peak timing for the later part of the EMREO waveform. The first peak after saccade offset was identified and its latency with respect to saccade onset was computed (y-axis) and expressed as a function of saccade duration (x-axis). Target locations are coded by color and each subject is identified with a unique symbol. By definition, the latency of this peak will scale with saccade duration, but the relationship could have been different for different subjects. Instead, we see that the relationship between saccade duration and the latencies of peaks clusters well around a single regression line for all subjects. (b) Similar to panel (a) but plots the latency of the first peak after saccade onset as a function of saccade duration. As before, saccades to each target location cluster together (by color) but the latency of this peak is unrelated to saccade duration.

In contrast, examining the relationship between saccade duration and the first peak of the EMREO after saccade onset for each subject and target, averaged across trials, indicates there is no apparent relationship between saccade duration and peak latency for this EMREO time window (Figure 8b). Again, the clustering by color indicates saccade duration is determined by target location. However, the peak latency does not change with changes in saccade duration. There is little evidence that peak *timing* during this temporal epoch covaries with target location: there is no obvious systematic color pattern along the x-axis for either the peaks or the troughs.

### Candidate metrics for future comparisons of EMREOs between normal and clinical populations: considerations for comparing across study designs and with limited data collection

One of the overarching motivations of this study is to identify the aspects of the EMREO that show the most stability across individuals with normal hearing, in anticipation of future comparisons to individuals with hearing loss. Studies in such clinical populations may be constrained in terms of data collection, requiring shorter testing sessions than are commonly used in research settings. What aspects of the experimental design can be sacrificed with minimal impact on the metrics being assessed? If necessary, testing could be limited to two horizontal targets, one on each side, such that there are contralaterally directed saccades for each ear. As shown in Figure 7a, the first 0-35 ms of the saccade-onset EMREO waveform varies with saccade amplitude but the amount of this variation is comparatively small. Figure 9 captures this in another way – here, we plot the peak-trough amplitude during the first 0-35 ms after saccade onset, averaged across subjects and plotted in different colors for the different target locations. Similar to previously published findings (Gruters et al., 2018), there is a small but reliable increase in peak-trough amplitude with increasing saccade amplitude. However, the impact of the change in saccade amplitude is negligible when averaging across all targets (gray bar, small standard error bars). This suggests that comparisons across studies that have used different horizontal target locations should be possible. Future work in clinical populations could rely on fewer targets, increasing statistical power for analysis at the single target level without increasing testing time.

**Figure 9.**
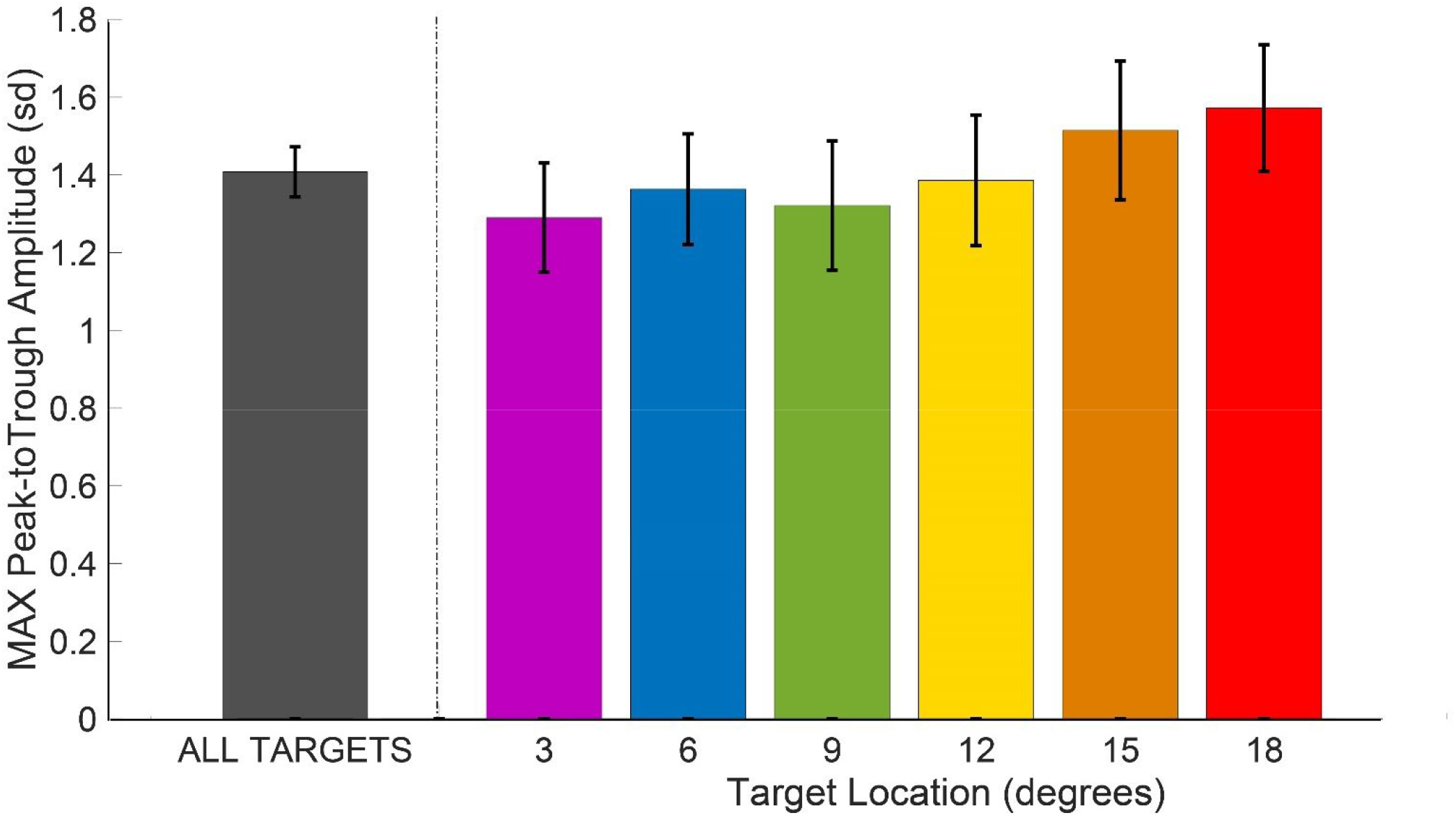
Average peak-trough amplitude scales with target location (colored bars; peaks and troughs obtained within 0-35 ms window after saccade onset for contralateral saccades; see also the population grand-average waveforms from *Figure 7a*). However, the range is narrow, such that pooling across locations loses little information: the gray bar for the 22 normal hearing subjects (44 ears) tested represents the population mean and standard error of the peak-to-trough amplitude. In short, the peak-trough amplitude 0-35 ms after onset of a contraversive saccade provides a good metric for EMREO amplitude, regardless of target location, providing a means for comparison across studies using different target locations and permitting use of a single contralateral target location if testing time/trial counts is a limiting factor in future studies.

Another choice to be made is how much data to collect to achieve reproducibility. To shed light on this question, we analyzed the correlations between the waveforms recorded in individual subjects across different blocks, focusing in on the 0-35 ms time window that is largely consistent across target locations, and pooling across those target locations to obtain an average contra-and an average ipsi-lateral waveform for each block for each subject (recall that subjects were tested in 4-11 blocks across 1-3 days; see Table 1 for details). We then computed the correlations between these average waveforms across blocks. Figure 10 shows the results for each subject, with the arrows/boundaries across days for subjects who were tested on more than one day. For contralateral saccades, the correlations between nearly all blocks in a given subject were strongly positive. The correlation patterns for ipsilaterally-guided saccades were also usually positive, but more exceptions emerged, suggesting that more testing is sometimes needed to accurately ascertain the ipsilateral waveform.

**Figure 10.**
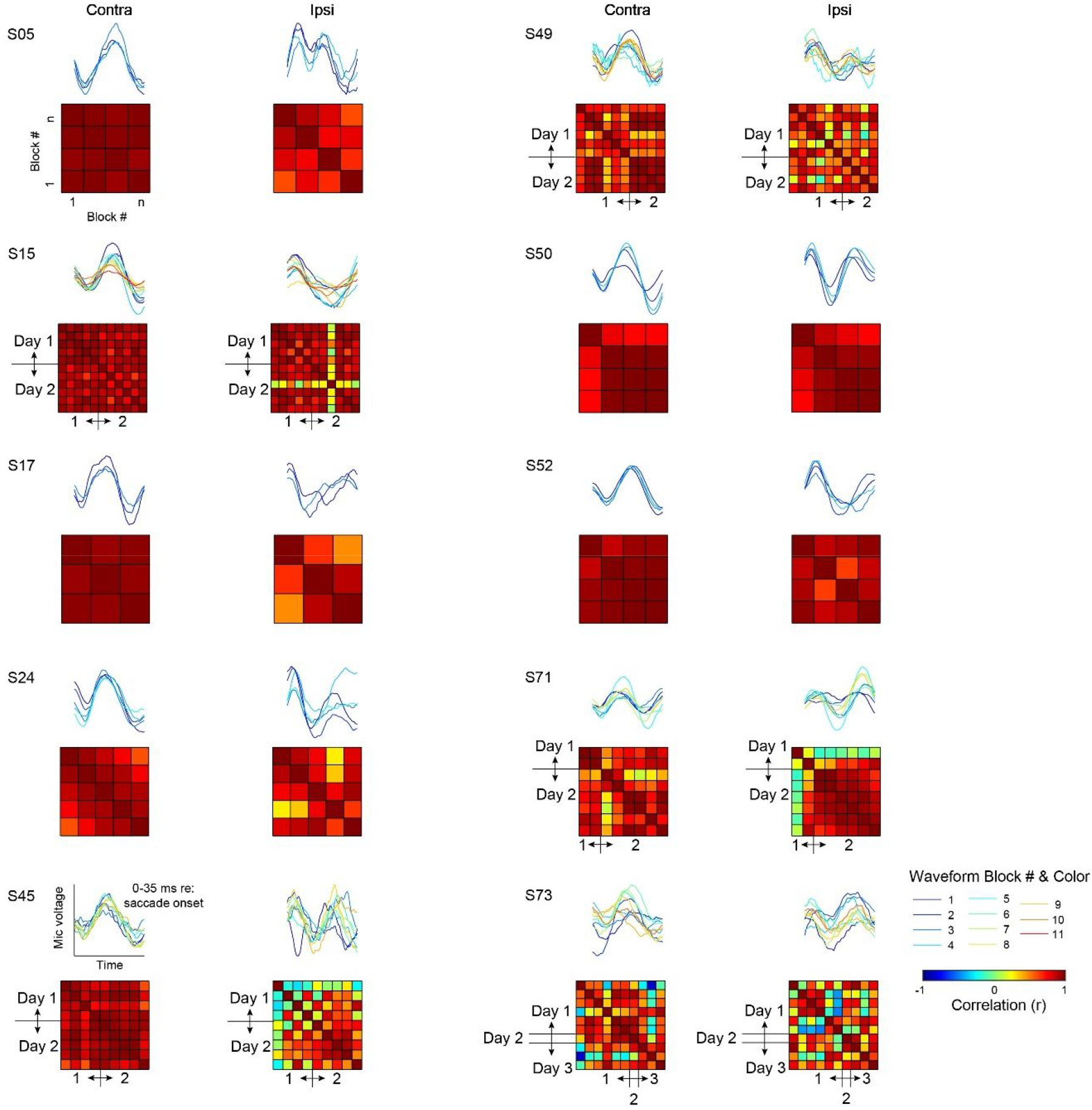
Reproducibility of EMREO waveforms across blocks and recording days in individual subjects. For each subject and recording block, an average EMREO waveform was computed for the 0-35 ms window in which the EMREO waveform was relatively insensitive to target location, permitting pooling across targets (as shown in *Figure 9*). The top panels show these average waveforms within each block. The lower panels show the correlations between these average waveforms across blocks.

On the whole, the amount of testing needed to obtain a reproducible results is modest. Each block yielded an average of 4 trials per target, or about 24 trials for each of the contralateral and ipsilateral hemifields (values computed from Table 1). Thus, in general, the minimum necessary amount of data to obtain a reproducible EMREO signal can be collected in on the order of ∼5 minutes of testing.

The bottom line of this and the preceding figures is that several metrics may prove useful for future clinical comparisons. Potential anomalies in EMREO “size” are likely to be best assessed using saccades to contralateral targets, via the maximum peak-trough amplitude observed within 0-35 ms of saccade onset, before any differences related to saccade offset time for saccades of different amplitudes have a chance to impact the waveform. Potential anomalies in EMREO timing are likely to be best assessed by evaluating (a) the timing of the first peak within 0-30 ms after saccade onset, and/or (b) the relationship between saccade duration and the later parts of the EMREO waveform after saccade offset.

## Discussion

Our primary goal was to identify the similarities and differences in EMREOs observed in participants with *normal* auditory function. Recognizing both the common and unique response properties of EMREOs sets the stage for understanding their purpose in normal hearing and for exploring the anatomical mechanisms responsible for generating these surprising signals in the ear. Ultimately, determining the EMREOs’ underlying anatomical mechanism(s) and their relationship to particular attributes of the EMREO signal may permit use of EMREOs as biomarkers, facilitating diagnosis of different types of auditory dysfunction. This latter goal is in the distant future; characterization of the normative population is a preliminary, but essential, step.

The first goal was to ascertain whether the EMREO is present in all normal hearing individuals. We observed EMREOs in all 10 of the subjects who participated in the saccade tasks explored here, consistent with our ongoing work and recent results from other studies (Lovich et al., 2022; Abbasi et al., 2023; Bröhl and Kayser, 2023; Lovich et al., 2023); (Murphy et al., 2020). This indicates that EMREOs are consistently present in the normal hearing population, which in turn suggests that EMREOs reflect a normal part of hearing.

We then considered which aspects of the EMREO show more vs. less variability, both within and across subjects. We reasoned that EMREO components similar across subjects may provide clues to either the perceptual purpose that EMREOs contribute to, the underlying mechanism that causes them, or both. We found that responses to horizontal targets were more consistent across subjects than responses to vertical targets, consistent with a role in spatial localization of sounds in an eye-centered reference frame: it is known that localization of sounds can be slightly more accurate in the horizontal than the vertical dimension (Tabry et al., 2013); (Jay and Sparks, 1990) (Makous and Middlebrooks, 1990), so to see a comparable difference in the horizontal vs. vertical dimensions of the EMREO provides a striking parallel. Relatedly, another recent study from our group has demonstrated that reconstruction of target locations from EMREOs is more accurate in the horizontal dimension than the vertical one (Lovich et al., 2022).

More puzzling is our finding that EMREO responses to contralateral targets are more consistent across subjects than those to ipsilateral targets. The interpretation of this finding is presently unknown. It seems unlikely to relate to any particular perceptual attribute of hearing, as what is ipsilateral to one ear is simultaneously contralateral to the other, so any negative perceptual consequences associated with this variability in signal should in theory balance out, at least for individuals with normal hearing in both ears. We therefore suspect that this asymmetry might ultimately prove to be connected to some aspect of the underlying mechanisms that generate EMREOs. It will therefore be of interest to evaluate whether this aspect of the EMREO phenomenon differs in clinical populations that have dysfunction involving either middle ear muscles or cochlear outer hair cells.

Our current findings confirm and extend our previous report that EMREOs are time-locked to both saccade onset and saccade offset (Gruters et al., 2018). We focused on the responses to contralateral targets as involving the most reproducible (i.e. least variable) EMREOs (across subjects: Figure 6; within subjects: Figure 10), and identified a clear pattern associated with saccade length/saccade duration/target location. This indicates a close connection between EMREOs and the control of saccades by the brain. Saccades are high speed eye movements, accomplished by “flooring the gas pedal.” To move the eyes different distances, the brain adjusts how long it floors the gas pedal. Indeed, saccades of varying sizes all reach peak velocity at about the same time (Van Opstal and Van Gisbergen, 1987), such that the velocity profile for all saccades in a given direction is nearly the same for the first few milliseconds. This parallels the similarity between EMREOs evoked for different saccades during the first 30 ms of the signal. The relationship between EMREOs and saccade length and duration, then, provides a parallel to the format of saccade commands, lending circumstantial support to the likelihood that some form of corollary discharge signal related to the saccade is issued by the brain and sent to the ear.

An important confirmatory finding in our present study concerns the estimated sound pressure level of EMREOs. In our present work, we estimated the typical EMREO associated with an 18 degree contralateral saccade to be range between ∼45-67 dB SPL (peak equivalent) across subjects with a mean of about 58.7 dB SPL (peak equivalent). This value is computed based on measurements taken with a sound level meter (accuracy +/-1.5 dB) and accords well with the value of 57 dB that we estimated previously (Gruters et al., 2018) based on a model of the ER10B+ microphone’s frequency response function in this low frequency range (Christensen et al., 2015).

A major goal for future work is to compare subjects with different types of auditory dysfunction to population data from normal hearing subjects. Studying patients with dysfunction involving any of the three types of actuators in the ear, i.e. the two middle ear muscles and the outer hair cells, will shed light on the contributions of these motor elements to EMREO generation. Different patterns of anomalies in the EMREO may occur in patients with different types of dysfunction. Studies in such clinical populations can be facilitated by focusing on analysis of EMREO components that exhibit the most consistency in subjects with normal hearing. Best practices can include focusing in on a single horizontal target in the contralateral and ipsilateral hemi-field and assessing the waveforms associated with these individual targets. If sampling across a range of locations in the horizontal or vertical dimensions is desired or if sampling is variable across different subjects, the regression model offers an ideal method of comparing across subjects. Our recent work using regression to compare across humans and monkeys performing a free-viewing task is a case in point (Lovich et al., 2023). Methods of ascertaining statistical significance with time-varying regression coefficients remain somewhat ad hoc, but the benchmarks used here (a criterion percentage of points differing from 0 in the -5-70 ms window around saccade onset) can provide a place to start. Cluster-based permutation methods such as those used in the EEG or fMRI literature will be needed for borderline cases (Nichols and Holmes, 2002; Maris and Oostenveld, 2007). Finally, the reproducibility of the average EMREO waveform across approximately 5 minute blocks of trials suggests that satisfactory EMREO assessment should be possible even in clinical settings where time of testing needs to be minimized.

Our overarching theory is that EMREOs are generated as a (by)product of processes involved in connecting the auditory scene to the visual scene across eye movements. Under this view, the underlying mechanisms associated with EMREOs may contribute to our experience of a perceptually stable world in which sights and sounds can be effortlessly linked to one another. Such linkage may involve implementing eye-movement-related adjustments to interaural timing and level differences via adjustments to the gain and timing of transmission through the middle ear, as suggested by recent work (Cho et al., 2023), which would help bring visual and auditory spatial signals into a common reference frame. The normative dataset presented here can provide a point of comparison not only for further study concerning those underlying mechanisms, but can also lead the way in investigations of what happens to our perception of an integrated visual-auditory scene when EMREOs deviate from the normal pattern.

Another future exciting area of research will be to identify the specific relationship between these peripheral signals and their associated causes/consequences at more central locations. Auditory cortex is known to exhibit visual sensitivity in both humans and monkeys (e.g. (Calvert et al., 1997) (Brosch et al., 2005; Ghazanfar et al., 2005) (Kayser et al., 2007)) ((for review, see for review, see Schmehl and Groh, 2021)), and signals related to eye movements have been identified as well (Werner-Reiss et al., 2003; Fu et al., 2004; Maier and Groh, 2010). Furthermore, several studies have identified an oscillatory change in excitability occurring time-locked to saccades and/or onsets of fixations (O’Connell et al., 2020; Leszczynski et al., 2023). While the sources of these latter effects are not well understood, the parallels to the influence of eye movements on the auditory periphery are striking.

## Acknowledgements

We would like to acknowledge former and current Groh lab members for thoughtful contributions throughout this research project: Jesse Herche, Bailey King, Dr. Jeff Mohl, Meredith Schmehl, Justine Shih, Chadbourne Smith, Chloe Weiser, Dr. Shawn Willett, and Tingan Zhu. We would also like to thank Hossein Abbasi, Dr. Patrick Bruns, and Dr. Jonathan Siegel for helpful discussions. This work was supported by NIH (NIDCD) grant DC017532.

## Footnotes

1 A key difference in this approach compared to fMRI general linear models (GLM) is that the time varying nature of the signal is captured via these fitted coefficients instead of pre-specified via so-called nuisance variables in a design matrix. proportion of p-values less than 0.05 was greater than the 5% that would be expected by random chance. We adopted a criterion of 10%, i.e. if more than 10% of these p-values were below 0.05, we considered the subject ear to have a statistically significant EMREO. This 10% threshold was adopted because a binomial test to evaluate whether one proportion is larger than another reaches a significance level of p<0.01 when considering 10% vs. 5% in a sample size of 149 values.

## Supplementary Materials

**Supplementary Figure 1.**
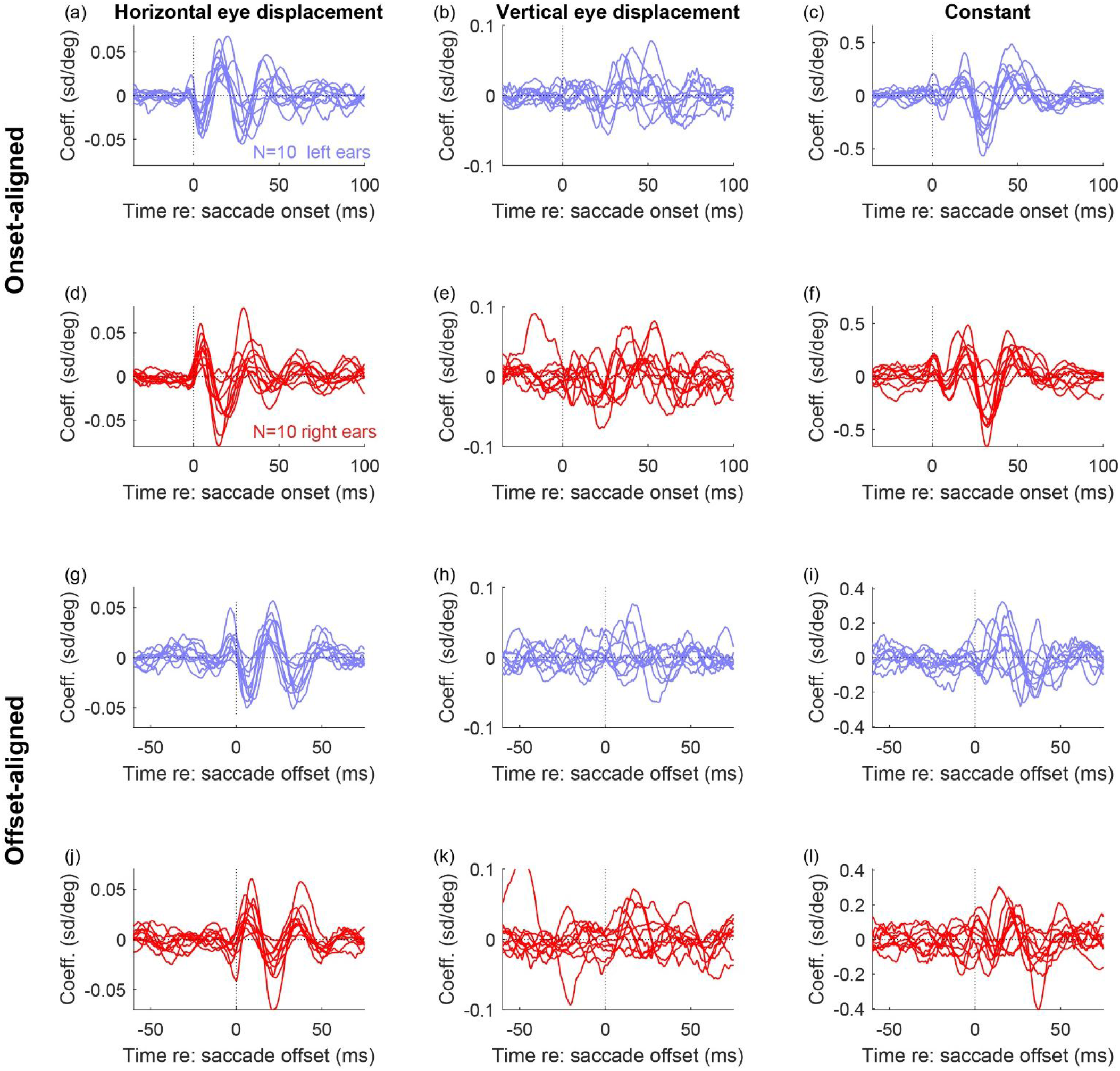
Regression curves for individual subjects (N=10) left ears (blue, a-c, g-i) and right ears (red, d-f, j-l), aligned on saccade onset (a-f) or offset (g-l). Same data as *Figures 2 and 3*.

